# glactools: a command-line toolset for the management of genotype likelihoods and allele counts

**DOI:** 10.1101/221127

**Authors:** Gabriel Renaud

**Affiliations:** Max Planck Institute For Evolutionary Anthropology, Deutscher Platz 6, Leipzig, Germany

**Author notes:** Present address: Center for GeoGenetics, Natural History Museum of Denmark, University of Copenhagen, 1350K Copenhagen, Denmark.

## Abstract

**Motivation:** Research projects involving population genomics routinely need to store genotyping information, population allele frequencies, combine files from different samples, query the data and export it to various formats. This is often done using bespoke in-house scripts which cannot be easily adapted to new projects and seldom constitute reproducible workflows.

**Results:** We introduce glactools, a set of command-line utilities which can import data from genotypes or population-wide allele frequencies into an intermediate representation, compute various operations on it and export the data to several file formats used by population genetics software. This intermediate format can take 2 forms, one to store per-individual genotype likelihoods and a second for allele counts from one or more individuals. glactools allows users to perform operations such as intersecting datasets, merging individuals into populations, creating subsets, perform queries (e.g. return sites where a given population does not share an allele with a second one) and compute summary statistics to answer biologically relevant questions.

**Availability:** glactools is freely available for use under the GPL. It requires a C++ compiler and the htslib library. (https://grenaud.github.io/glactools/).

**Supplementary information:** Supplementary methods and results are available at *Bioinformatics* online.

## 1 Introduction

Advances in next-generation sequencing have enabled researchers to produce large amounts of high-resolution genotype data for individuals pertaining to populations of interest. Advances in ancient DNA have also enabled the comparison of historical and contemporary samples. Publicly available datasets, such as high-resolution genomes (Abecasis *et al*. (2012); Mallick *et al*. (2016)), provide valuable resources when analyzing newly sequenced individuals. Software such as bcftools (Li *et al*. (2009)) enables users to store genotype data as VCF/BCF files, intersect datasets from different individuals and export them to different formats as well. More recent work has focused on how to efficiently index, query and store genotyping data (Li (2015); Layer *et al*. (2016); Zheng *et al*. (2017)).

Many different native file formats are used by programs aimed at population genetics analyses. Several of those do not incorporate genotype likelihoods into their statistical models but use instead allele frequencies or pseudo-haploid data in the form of single bases. Users are thus faced with the problem of exporting the data to such formats. Another example of a typical task involves intersecting this genotyping data with heterogeneous data such as population allele frequencies from third-party sources or ancestral allele information.

Research groups typically resolve these problems using custom inhouse scripts. In addition, such bespoke scripts are used to compute routine queries such as restricting analyses to transversions or to sites where the ancestral allele is present in a given population. However, such methodology is seldomly portable from project to project. Often enough, such workflows are completely irreproducible even within the same group and are usually rewritten once the original programmer leaves the group.

We present glactools, a set of programs aimed at exporting various formats (e.g. VCF, BAM, AXT) into compressed binary files and perform operations on those. We introduce 2 types of binary file formats, GLF for storing genotype likelihoods and ACF for allele counts. The GLF format contains genotype likelihoods for single individuals. The ACF format stores the number of times a specific base is observed in an individual or population. Both of these files formats contain the following genomic information: chromosome name, coordinate, reference and alternative allele. They also store sites that are homozygous for the reference base instead of simply storing variants as several program distinguish between the absence of a segregating site and missing data.

This data is encoded as block compressed binary files that can be indexed for retrieval of specific regions using the same methodology used by htslib from SAMtools (Li *et al*. (2009)). To provide a traceable workflow, each GLF/ACF file contains a header detailing the precise commands used by each subprogram to produce this file all the way to its creation from raw data. Ancestral allele information can be added from multiple sequence alignments to other species. The resulting GLF/ACF files can be united, intersected and filtered for certain criteria (e.g. transversions, presence of a derived allele in population X but not in Y). Summary statistics such as average coalescence (Prüfer *et al*. (2010)) or D-statistics (Patterson *et al*. (2012)) can be computed using glactools directly on those files. Multiple CPUs can be used to efficiently compute such statistics. Although our approach was originally designed to work with hominin species, our framework is easily adaptable to work with any type of autosomal data. Our open-source C++ framework also allows programmers to write custom applications and filters.

## 2 Methods

glactools represents genotype likelihoods or allele counts as block compressed binary files that can be indexed (see Figure 1). GLF contains genotype likelihoods from a single diploid individual and ACF can contain data for 1 or more individuals (see Supplementary Methods for further details). Unlike VCF/BCF, the GLF format merely stores the genotype likelihoods and assumes that the input data has been pre-filtered for quantities such as mapping quality, mappability, coverage etc. Both the ACF and GLF files do not currently support phasing, triallelic sites or indels as they are aimed at intra-species studies. This intermediate file format is efficient as both ACF/GLF can be upwards of 2X smaller in terms of file size compared to BCF/VCF files.

**Fig.1.**
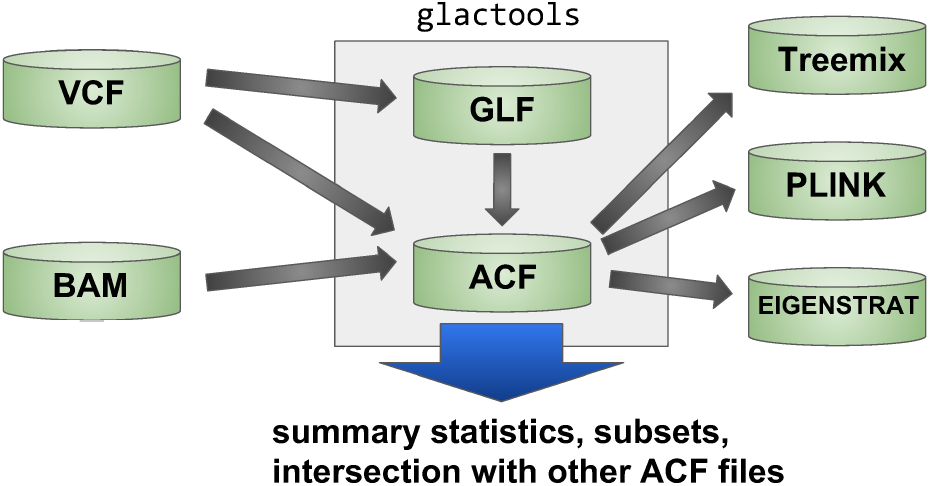
Schematic view of glactools’ capabilities.

A single line of ACF (see below) contains the information for an outgroup (root), the most recent common ancestor (anc) and the population in question (pop&#x0023;1). The allele count represents the number of reference and alternative alleles found at that given position and a binary flag indicating whether the site is a putative CpG site or not:

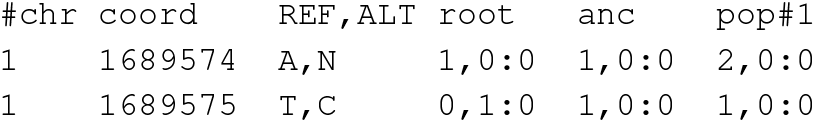

glactools allows data to be imported from VCF files, BAM files or 23 andMe files. glactools can import ACF files from VCF files or transform GLF files into ACF using likelihood ratios on the genotype likelihoods to infer the number of alleles present in the individual (see Supplementary Methods). The base count from BAM files can also be used for comparison between samples with extreme differences in coverage.

Once data has been imported, glactools can create intersections using multiple samples. Allele counts from various populations can be merged together. Users can filter for sites where two individuals share an allele or where they do not. Sharing or not sharing an allele with the ancestral state filters for a population with the ancestral or derived allele respectively. It is also possible to filter specific sites using a BED file, for example, to retain sites within regions of interest. Summary statistics such as all pairwise average coalescence, site frequency spectrum and D-statistics can also be computed from the allele count matrix. Finally, data can be exported to either FASTA, NEXUS, PLINK, EIGENSTRAT, G-PhoCS or TreeMix.

## 3 Results

glactools has various commands to import, transform and export data. We summarize certain key commands in Table 1. Documentation for all commands is included with the package.

**Table 1.** Examples of commands for glactools.

| command | input | output |
| --- | --- | --- |
| view | GLF/ACF binary to/from text format |  |
| bam2acf | BAM | ACF |
| vcf2acf | VCF | ACF |
| vcf2glf | VCF | GLF |
| meld | ACF | ACF with merged populations |
| intersect | GLF/ACF | GLF/ACF intersection of input files |
| segsite | ACF | ACF with only segregating sites |
| glf2acf | GLF | ACF with called allele counts |
| acf2treemix | ACF | TreeMix input format |
| acf2EIGENSTRAT | ACF | Eigenstrat format |
| acf2bplink | ACF | PLINK format |

Commands that read, write and transform ACF/GLF files .

As an example of a potential analytical workflow with glactools, we transformed the Phase III data from the 1000 Genomes (Abecasis *et al*. (2012)) into TreeMix and ADMIXTURE input. The VCF files were transformed into an ACF file, alleles counts for individuals from the same population were merged and finally exported to TreeMix and ADMIXTURE input. The commands necessary to complete these steps are presented in Supplementary Results.

## Acknowledgements

We are indebted to Johannes Krause whose ancient DNA project provided the initial impetus for this project. We would also like to thank Ana T. Duggan, Ilan Gronau, Janet Kelso, Ludovic Orlando, Martin Sikora and Thorfinn Sand Korneliussen for helpful feedback.

## Funding

Work was funded by the Max Planck Society and NSERC for a PGS D scholarship and currently funded by a Marie Skłodowska-Curie IF.

